# A re-inducible gap gene cascade patterns the anterior-posterior axis of insects in a threshold-free fashion

**DOI:** 10.1101/321786

**Authors:** Alena Boos, Jutta Distler, Heike Rudolf, Martin Klingler, Ezzat El-Sherif

## Abstract

Gap genes mediate the division of the anterior-posterior axis of insects into different fates through regulating downstream hox genes. Decades of tinkering the segmentation gene network of the long-germ fruit fly *Drosophila melanogaster* led to the conclusion that gap genes are regulated (at least initially) through a threshold-based French Flag model, guided by both anteriorly- and posteriorly-localized morphogen gradients. In this paper, we show that the expression patterns of gap genes in the intermediate-germ beetle *Tribolium castaneum* are mediated by a threshold-free ‘Speed Regulation’ mechanism, in which the speed of a genetic cascade of gap genes is regulated by a posterior gradient of the transcription factor Caudal. We show this by re-inducing the leading gap gene (namely, *hunchback*) resulting in the re-induction of the gap gene cascade at arbitrary points in time. This demonstrates that the gap gene network is self-regulatory and is primarily under the control of a posterior speed regulator in *Tribolium* and possibly all insects.

## Introduction

The French Flag model is one of the earliest models of pattern formation in development (1), in which thresholds of a morphogen gradient (e.g. T1 and T2 in Figure 1 A’) set the boundaries between different gene expression domains. Recent studies of morphogen-mediated patterning, however, presented several challenges to this simple picture. First, gene expression domains are usually found to be dynamic, and in many cases, are expressed sequentially in a wave-like fashion (e.g. during neural tube and limb bud patterning in vertebrates and during anterior-posterior (AP) fate specification in vertebrates and insects) (2–12). Even if activated simultaneously, gene expression domains usually undergo continuous shifts in space (e.g. gap and pair-rule domains in *Drosophila*) (13–15). In such cases, it is difficult to correlate the locations of gene expression domains with specific values of morphogen concentrations. Second, morphogen exposure time was found to have a crucial effect on patterning. For example, the exposure time of Caudal (Cad) regulates the timing of gap and pair-rule genes in insects and that of Hox genes in vertebrates (11,16). Similarly, the concentration and exposure time of Sonic hedgehog (Shh) determines which fate a cell in the vertebrate neural tube and limb bud will take (6–8).

**Figure 1.**
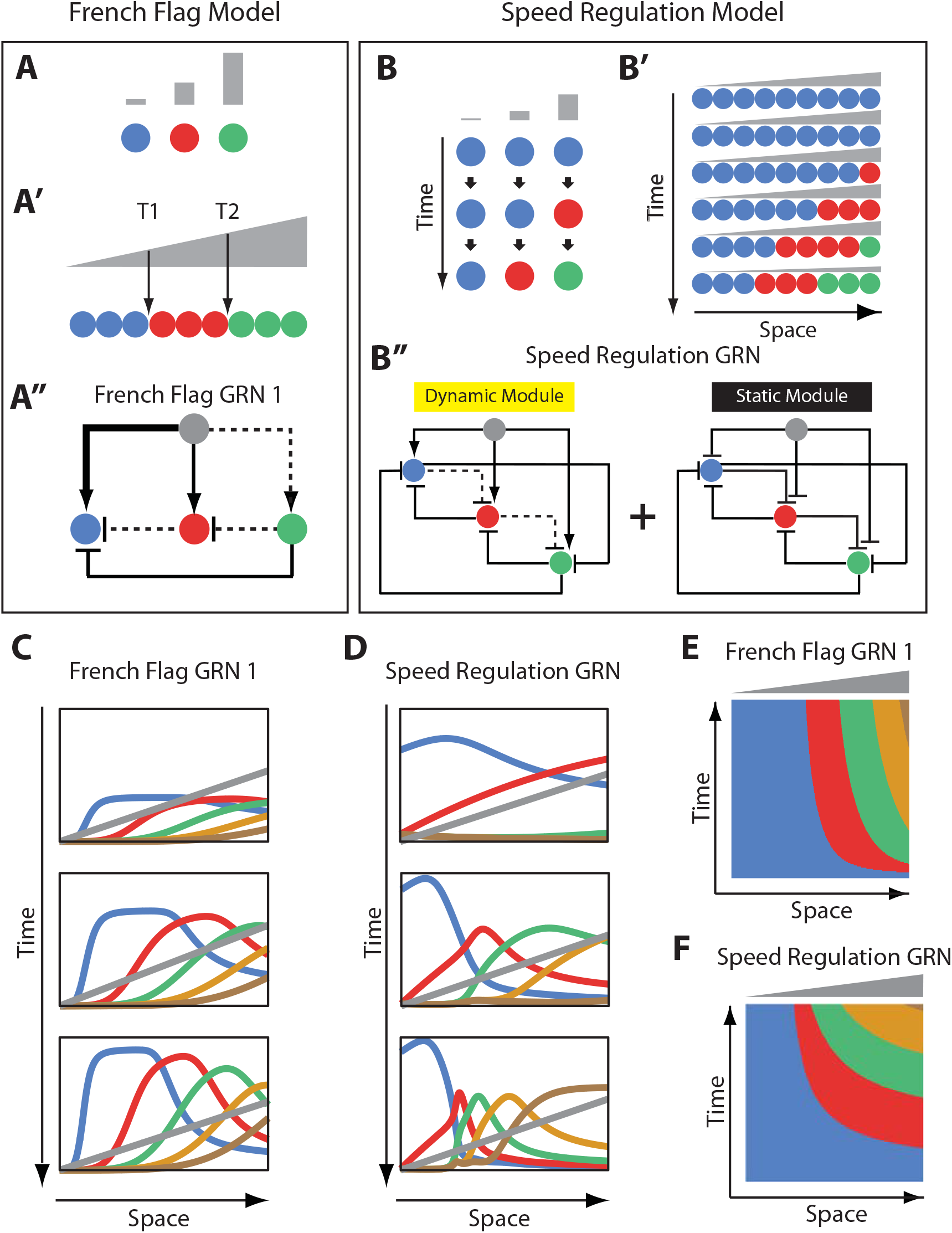
French Flag model versus Speed Regulation model. (**A-A”**) In the French Flag (FF) model, different concentrations of a morphogen gradient (grey) activate different cellular states (**A, A”**) based on a set of morphogen thresholds (here T1 and T2; **A’**). In **A** and **A’**, cells are represented by circles and cellular states are shown in blue, red and green. Shown in **A”** is a GRN realization of the FF model (GRN 1; genes representing different cellular states are shown in circles; arrowheads stand for activation, and flat bars stand for repression; the thicker the line, the stronger the activation/repression; dashed lines stand for weak activation/repression). (**B-B”**) In the Speed Regulation (SR) model, all cells (shown in circles in **B** and **B’**) transit through different cellular states (shown in blue, red, and green) with a speed that is proportional to the concentration of a morphogen gradient (grey). Shown in **B”** is a GRN realization of the SR model (genes representing different cellular states are shown in circles; arrowheads stand for activation, and flat bars stand for repression; dashed lines stand for weak activation/repression). (**C,E**) Computer simulation of FF GRN 1 shown as plots of expression domains along space for selected time points (**C**) and as a kymograph (**E**). (**D,F**) Computer simulation of SR GRN shown as plots of expression domains along space for selected time points (**D**) and as a kymograph (**F**).

For these reasons, recent works in morphogen-mediated patterning are suggesting a more dynamic and time-based paradigm rather than threshold-based (2,6,11,14,15,17–23). However, the general description of the French Flag model (1) (Figure 1 A, A’) is purely phenomenological (i.e. descriptive). Hence, it is unclear if a gene regulatory network (GRN) realization of the French Flag model, while still threshold-based, would reconciliate the experimentally observed deviations (namely, the dynamic nature of gene expression domains and the sensitivity of patterning to morphogen exposure times). In this paper, we show that indeed important classes of GRN realization of the French Flag model “transiently” exhibit exactly these features. In particular, at an initial transient phase, gene expression domains are dynamic and keep shifting and shrinking as long as the morphogen gradient is applied, hence its sensitivity to the exposure time of the morphogen gradient. However, gene expression domains finally stabilize at the thresholds set by the morphogen gradient, adhering to the tenets of the French Flag model. Hence, we argue that the defining feature of the French Flag model is not its transient dynamics, but its threshold-based steady state behavior.

However, a recently devised mechanism (termed ‘Speed Regulation’ model, Figure 1 B, B’) (11,12), that has been suggested to be the basis of the AP fate specification in insects, is purely time-based and threshold-free. In this model, segmentation genes of the gap class are wired into a genetic cascade, so that gap genes are activated sequentially in time. The temporal progression of gap gene expressions is translated into a spatial pattern through modulating the speed (or timing) of the gap gene cascade by a morphogen gradient of the transcription factor *caudal* (*cad*).

In contrast to the French Flag model, the Speed Regulation model is completely threshold-free, where gene expression domains keep shifting and shrinking as long as the morphogen gradient is applied (without ever reaching a steady state) until the gradient is retracted. In this paper, to demonstrate the threshold-free nature of gap gene regulation, we re-induce the leading gene in the gap gene cascade (namely, *hunchback, hb*) at arbitrary times during the AP specification phase of the beetle *Tribolium castaneum* using a transgenic line carrying a heat-shock-driven *hb* CDS. This resulted in resetting the gap gene cascade and re-establishing the gap gene expression sequence in time and space. This argues against any spatial or temporal signal that sets the locations of gap gene expression boundaries in a threshold-based fashion. Using computational modeling, we show that this self-regulatory behavior of gap gene regulation is difficult to explain using different realizations of the French Flag model and, alternatively, is a hallmark of time-based (or ‘speed regulation’-based) patterning.

The paper is organized as follows. First, we contrast the French Flag model to Speed Regulation model and their corresponding GRN realizations, highlighting their similarities and differences. We argue that the basic French Flag model, in contrast to the Speed Regulation model, is incapable of reproducing the phenomenology of insect development and evolution. Secondly, we use a modified version of the French Flag model (called ‘French Flag with a Timer Gene’) as a possible contender to the Speed Regulation model in explaining the AP axis fate specification in insects. We show that the major difference between the two models is that the French Flag model (and its modified version) depends on morphogen thresholds in setting the boundaries between gene expression domains, whereas the Speed Regulation model is self-regulatory and threshold-free. We suggest an experimental test to differentiate between the two mechanisms: whether the patterning scheme can be reset independently from any temporal or positional signal. We then carry out this test experimentally in the beetle *Tribolium castaneum*, concluding that the Speed Regulation model is a more plausible mechanism to explain insect development and evolution.

## Results

### Comparing French Flag and Speed Regulation models in patterning non-growing tissues

A common problem in development is how to divide a group of cells into different identities, each specified by the expression of one or more genes. Here we present two different patterning mechanisms to partition a static (i.e. non-growing) tissue along a spatial axis: the French Flag (FF) model (Figure 1 A, A’) (1) and the Speed Regulation (SR) model (Figure 1 B, B’) (11,12). In the FF model, different ranges of a morphogen concentration (grey in Figure 1A) activate different cellular states (specified by the expression of one or a group of genes; different states are given different colors in Figure 1A). Figure 1A’ shows how this scheme can be used to partition a row of cells into different cellular states using a gradient of the morphogen along space, where morphogen concentrations above zero and below threshold T1 activate the blue state, above T1 but below T2 activate the red state, and above T2 activate the green state.

In the SR model, all cells have the capacity to transit through all cellular states in the same order (blue, red, then green in Figure 1B). However, in contrast to the FF model, different morphogen concentrations do not directly activate specific cellular states but regulate the *speed* of state transitions in time (Figure 1B). Applying a gradient of the morphogen along a row of such cells induces kinematic waves of cellular states that propagate from high to low values of the gradient (Figure 1B’). Eventually, cells are partitioned into different domains of cellular states, and the morphogen gradient should decay and/or retract to stabilize the pattern (otherwise the expression pattern would continue to shrink and propagate towards the lower end of the gradient; last row of cells in Figure 1B’).

While the final results of both models are the same, their dynamics look very different. Whereas bands of cellular states in the FF model are established simultaneously without any need for a temporal component, cellular states are very dynamic in the SR model, where timing is very critical (both morphogen concentration and exposure time determine which state a cell will end up in). However, the absence of the temporal component in the French Flag model is probably unrealistic since any real life molecular implementation of the model would exhibit some transient dynamics. Hence, we aim here to use GRN realizations for both the FF and SR models and compare them on equal footing.

Using *in silico* evolution techniques, Francois and Siggia in ref (24) performed an unbiased exploration of possible morphogen-regulated GRNs that can divide an embryonic tissue into different fates. Although several solutions were found, they were mostly variations on the same underlying principle, which happened to be a straightforward realization of the FF model. A simple instance of this family of GRN solutions is shown in Figure 1A” using 3 genes (more genes can be added to the scheme in a straightforward manner; see Text S1). In the example GRN in Figure 1A”, the morphogen gradient (grey) activates different genes with different strengths: strongly activates the blue gene, moderately activates the red gene, and weakly activates the green gene. Being strongly activated by the morphogen gradient, the blue gene is first activated in all cells exposed to the morphogen gradient. Being moderately activated by the morphogen, the red gene is then (after a delay) activated only at regions where the morphogen value exceeds its activation threshold. Since the red gene represses the blue gene, the blue gene now turns off where the red gene is now expressed. In a similar fashion, the green gene turns on (after a delay) only in regions where the morphogen concentration exceeds its activation threshold and then, eventually, represses the red gene there (see simulation of a 5-genes FF GRN in Figure 1C and Movie S1). This and conceptually similar GRN realizations of the FF model (which we will call ‘French Flag GRN 1’ or ‘FF GRN 1’; Figure 1A”) encode morphogen activation thresholds ‘explicitly’, where different genes have different binding affinities to the morphogen molecules. However, other realizations could possess ‘implicit’ thresholds encoded by the overall connectivity of the GRN. An example for this family of networks is the AC-DC network motif (which we will call here ‘French Flag GRN 2’ or ‘FF GRN 2’; Figure S1; Movie S2) suggested as the underlying mechanism for patterning the vertebrate neural tube (3). Although the structure and logic of FF GRNs 1 and 2 are different, they both exhibit similar transient dynamics and eventually reach specific steady states set by morphogen thresholds, either explicitly or implicitly (compare Movies S1 and S2).

Now we turn to a molecular realization for the SR model recently suggested in refs (11,12). In this realization, two GRN modules are employed: a dynamic module and a static module (see Figure 1B” for a 3-genes realization and Text S1 for a 5-genes realization). The dynamic module is a genetic cascade that mediates the sequential activation of its constituent genes. The static module is a multi-stable network that mediates the refinement and stabilization of gene expression patterns. The morphogen gradient activates the dynamic but represses the static module. Hence, as we go from high to low values of the morphogen gradient, the dynamic module experiences excessively higher stabilizing effect from the static module, and consequently, runs slower. This is a straightforward realization of the core mechanism of the SR model (Figure 1 B, B’) and, hence, a morphogen gradient applied to such scheme induces sequential kinematic waves that propagate from the high to the low end of the gradient as shown by the computer simulations in Figure 1D and Movie S3.

Comparing the spatiotemporal dynamics of the molecular realizations of the FF model (both GRN 1 and GRN 2) and that of the SR model shows striking similarities, where gene expression domains are activated sequentially at the high end of the morphogen gradient (grey) and propagate in kinematic waves towards the low end of the gradient (compare Figure 1C,E to 1D,F and Movie S1 to S3). Both morphogen concentration and exposure time are important factors to determine which cellular state a certain cell will have at a certain point of time (at least at the initial transient phase). A main difference, however, is that gene expression domains in the FF model realizations are only dynamic during the initial transient phase, but eventually reach a steady state where they stabilize at certain morphogen thresholds (Figure S2 A, Movies S1 and S2). These steady state thresholds are set explicitly, as in FF GRN 1, or implicitly, as in FF GRN 2. On the other hand, gene expression domains in the SR model keep shrinking and propagating towards the low end of the morphogen and never stabilize or reach a steady state (Figure S1 C, Movie S3), unless the morphogen gradient decays or retracts (Movie S4).

### Comparing French Flag and Speed Regulation models in reproducing the phenomenology of insect development and evolution

So far, we have considered the application of the FF and SR models in patterning a static group of cells (i.e. a non-growing tissue). Here we consider their application to the problem of insect development and evolution, where a patterning mechanism is needed to pattern both growing and non-growing tissues.

The anterior fates of insects (Figure 2A) form in a non-growing tissue (called the ‘blastoderm’), whereas posterior fates form in a growing tissue (called the ‘germband’) (25,26). The number of fates specified in the blastoderm versus germband differ in different insects. In short-germ insects, (most of) AP fates are specified in the germband (first column in Figure 2A). In long-germ insect, (most of) AP fates form in the blastoderm before transiting into the germband stage (third column in Figure 2A). Intermediate-germ insects lie somewhere in between these two extreme cases, where several fates are specified in the blastoderm, and the rest in the germband (second column in Figure 2A). Throughout evolution, the specification of AP fates seems to shift easily from the germband to the blastoderm, resulting in a trend of short-germ to long-germ evolution (25,27,28) (with some reports of the opposite evolutionary path (29)). This hints that AP fates in both blastoderm and germband are likely to be specified using the same mechanism. Indeed, we have recently shown that in the intermediate-germ insect *Tribolium castaneum*, blastodermal fates can form in the germband, and germband fates can form in a non-growing (blastoderm-like) tissue (11).

**Figure 2.**
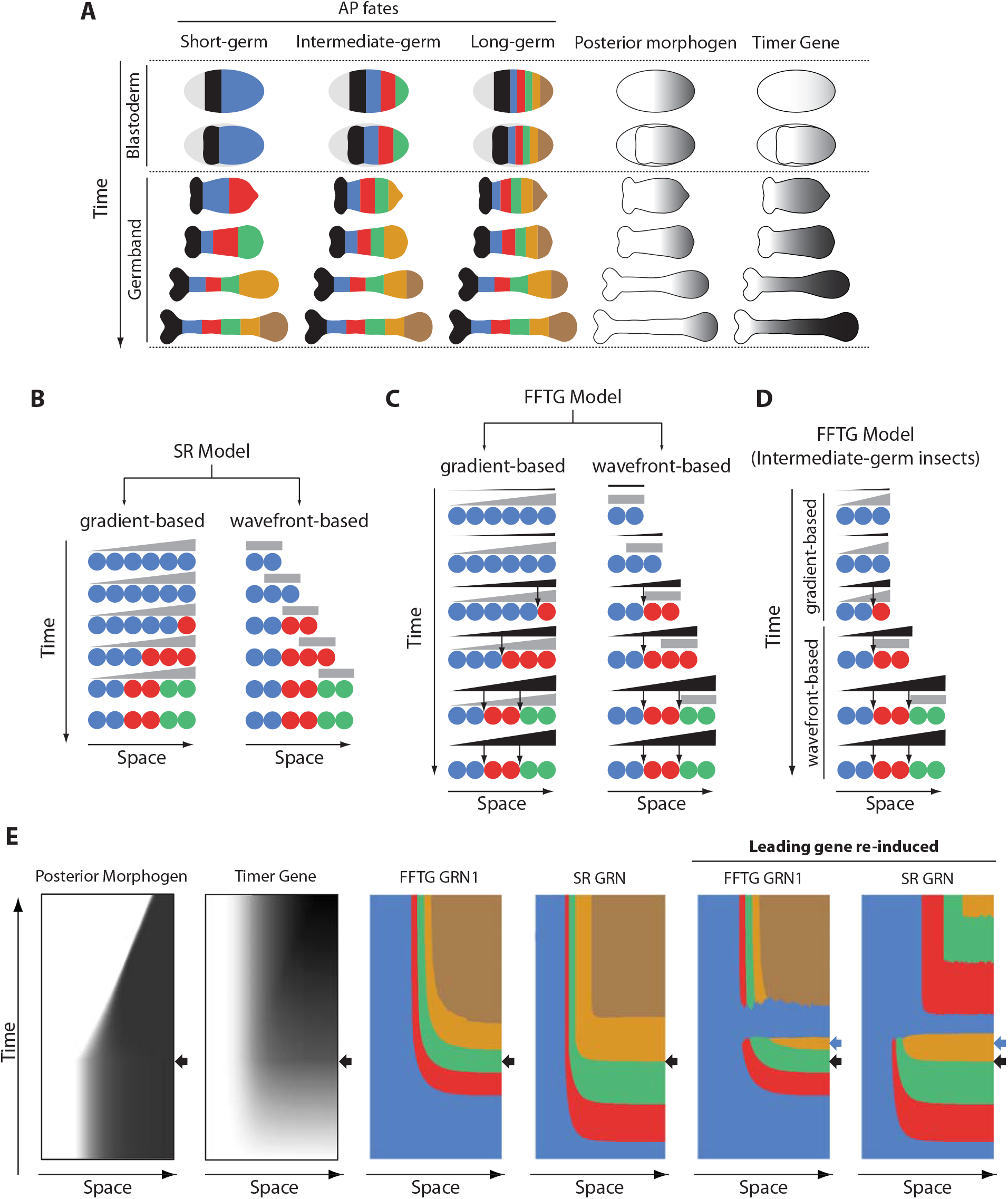
Application of FF and SR model to the problem of insect development and evolution. (**A**) Anterior-posterior (AP) fate specification in insects of different germ types: short-, intermediate- and long-germ. Different fates are shown in different colors. In short-germ embryogenesis, most of fates are specified in an elongating embryonic structure called ‘germband’. In long-germ embryongenesis, most of fates are specified in a non-elongating embryonic structure called ‘blastoderm’. In intermediate-germ embryogenesis, anterior fates are specified in the blastoderm, while posterior fates are specified in the germband. A posteriorly-localized morphogen is shown in grey. Timer Gene, as required by French Flag with Timer Gene (FFTG) model is shown in black with different shades (the darker the higher the concentration). (**B**) The Speed Regulation (SR) model can operate in two modes: gradient-based and wavefront-based. In the gradient-based mode, a static gradient of the speed regulator (grey) is applied to a static field of cells (shown in circles). In the wavefront-based mode, a boundary of the speed regulator retracts along an elongating field of cells. (**C**) The FFTG model can operate in a gradient-based or a wavefront-based mode as well. The Timer Gene (black) is activated by a posterior morphogen (grey) and suffers little or no decay so that it unfolds into a long-range gradient along the entire AP axis in either the gradient-based or the wavefront-based mode. Different concentrations of the Timer Genes (arrows) set the boundaries between different fates. (**D**) FFTG model can pattern the AP axis of intermediate-germ insects: acting in the gradient-based mode to pattern anterior fates (during the blastoderm stage) and acting the wavefront-based mode to pattern posterior fates (during the germband stage). (**E**) Kymographs of FFTG and SR GRNs when applied to AP patterning of intermediate-germ insects (first 4 panels; black arrow marks blastoderm-to-germband transition). Panels 5 and 6 show the response of the FFTG and SR GRNs to the re-activation of the leading gene (blue; the time of blue gene re-induction is marked by a blow arrow). In FFTG GRN, normal expression pattern is restored after a brief dominance of the blue gene. In SR GRN, already formed expression is down-regulated and the genetic cascade is reset.

In the derived AP patterning of *Drosophila* (and potentially all other dipterans), AP fate specification is mediated by an anterior morphogen (*bicoid* (*bcd*)) while *cad* serves as additional activator of posterior gap genes (26,30). However, most insects lack *bcd* (31,32), and AP fates might rely solely on a posterior morphogen (Figure 2A, fourth column), as suggested by recent works (9,11,20). In ref (11), it was suggested that AP fate specification in short- and intermediate-germ insects is mediated by the SR model, where a posterior morphogen (*cad* in *Tribolium*) regulates the sequential activation of the AP-determinant genes of the gap class. This suggestion was based on the observation that the SR model can operate in two modes (Figure 2B): a gradient-based, which can pattern a non-elongating tissue (as discussed in the previous section), and a wavefront-based mode in which the posterior morphogen gradient continuously retracts as the tissue elongates, in a set-up similar to the Clock-and-Wavefront model (33–36). The wavefront-based mode is best suited for patterning elongating tissues. It was also shown that the flexibility of the SR model to pattern both elongating and non-elongating tissue could offer an evolutionary mechanism where a smooth transition between short- and long-germ modes of insect development is possible (Movie S5).

However, in ref (24), Francois and Siggia suggested a simple modification that enables also the FF model to exhibit such flexibility in patterning both, elongating and non-elongating tissues. In this scheme, the posterior morphogen (fourth column in Figure 2A, and grey in Figure 2C) activates a gene (termed a ‘Timer Gene’ (24); fifth column in Figure 2A, and black in Figure 2C). The Timer Gene is assumed to have negligible decay rate, so that it continuously builds up in a non-growing tissue (left column in Figure 2C). In a growing tissue, the expression of the Timer Gene (black in Figure 2C, right column) builds up in the presence of the retracting posterior gradient (grey in Figure 2C, right column), while it stabilizes upon its retraction. Hence, a long-range gradient of the Timer Gene forms along the whole axis of the full-grown tissue (last row in the fifth column of Figure 2A and the right column of Figure 2C). Thresholds of different concentrations of the Timer Gene then set the boundaries between different fates, in a similar fashion to the FF model (thresholds are shown in arrows in Figure 2C). We call this scheme ‘French Flag model with a Timer Gene’ (FFTG). We notice that both the SR model and the FFTG model can operate in gradient-based and wavefront-based modes (and, hence, can pattern both non-elongating and elongating tissues) and exhibit similar dynamics and final pattern (compare Figure 2B and 2C; see Figure S2).

A transition from a gradient-based to wavefront-based patterning would still result in a long-range gradient of the Timer Gene along the tissue axis (Figure 2D). Hence, similar to the SR model, the FFTG model can mediate AP fate specification in insects of different germ types: operating in the wavefront-based mode for short-germ insects, operating in the gradient-based mode for long-germ insects, and operating in the gradient-based then switching to wavefront-based mode for intermediate-germ insects (Movies S6; compare to Movie S5).

Since gene expression dynamics of both the SR and FFTG models are very similar, we sought an experimental test to differentiate between the two models. In our demonstration of the experimental test, we will use the GRN realization of both SR and the FFTG models (5-genes version of Figures 1B” and 5-genes version of 1A’’ after adding a Timer Gene, respectively; see Text S1). We apply both models to the case of AP fate specification of an intermediate-germ insect.

Here we note that the sequential activation of genes in the SR model is mediated by the interaction between fate-specifying genes themselves, whereas the Timer Gene is the mediator of sequential gene activation in the FFTG model. Hence, force-resetting the fate-specification gene sequence will reset the SR model, whereas resetting the expression pattern for the FFTG model would require resetting the Timer Gene instead. This is evident in our simulations of both models in Figure 2E and Movies S7 and S8. After the expression of the first three AP-specifying genes (blue, red, and green in Figure 2E), the blue gene was briefly re-induced. In the FFTG model, the blue gene becomes briefly dominant, but the already formed pattern and the newly forming pattern are largely unchanged (compare 5^th^ to 3^rd^ column of Figure 2E). This is a natural consequence of the fact that the Timer Gene (Figure 2E, second column) is the main driver of the patterning process, which is unaffected by the blue gene re-induction. On the other hand, re-inducing the blue gene had two consequences in the case of our GRN realization of the SR model (compare 4^th^ and 6^th^ columns of Figure 2E): (*i*) the already formed pattern outside of the expression domain of the posterior morphogen is deleted and dominated by the continued expression of the blue gene, and (*ii*) the temporal gene sequence is re-established within the expression of the posterior morphogen, resulting in the re-establishment of the patterning process. This dual effect results from the dual regulation mode of our realization of the SR model: a dynamic genetic module is active within the posterior morphogen expression domain, whereas a static module is active outside. In our GRN realization, re-inducing the blue gene resets the dynamic module (which is basically a genetic cascade), while it down-regulates all the genes of the static module except the re-induced blue gene (since it is a multi-stable mutually exclusive GRN).

### A dual response upon re-inducing *hunchback* in the *Tribolium* embryo

During AP patterning of the *Tribolium* embryo, gap genes (namely, *hunchback* (hb), *Krüppel* (*Kr*), milles-pattes (*mlpt*), and *giant* (*gt*); Figure 3) (37–42) are expressed in sequential waves of gene expressions that propagate from posterior to anterior in the presence of a gradient of the master regulator *caudal (cad)* (11). Hereafter, we will call the region of the embryo where *cad* is expressed: the ‘active-zone’. Upon retraction of the *cad* gradient, gap gene expressions stabilize into static domains before they gradually fade. Some of the gap genes have two trunk expression domains, namely: *hb, mlpt* and *gt* (shown in blue, green and gold, respectively in Figure 3; late trunk domains are outlined in black).

**Figure 3.**
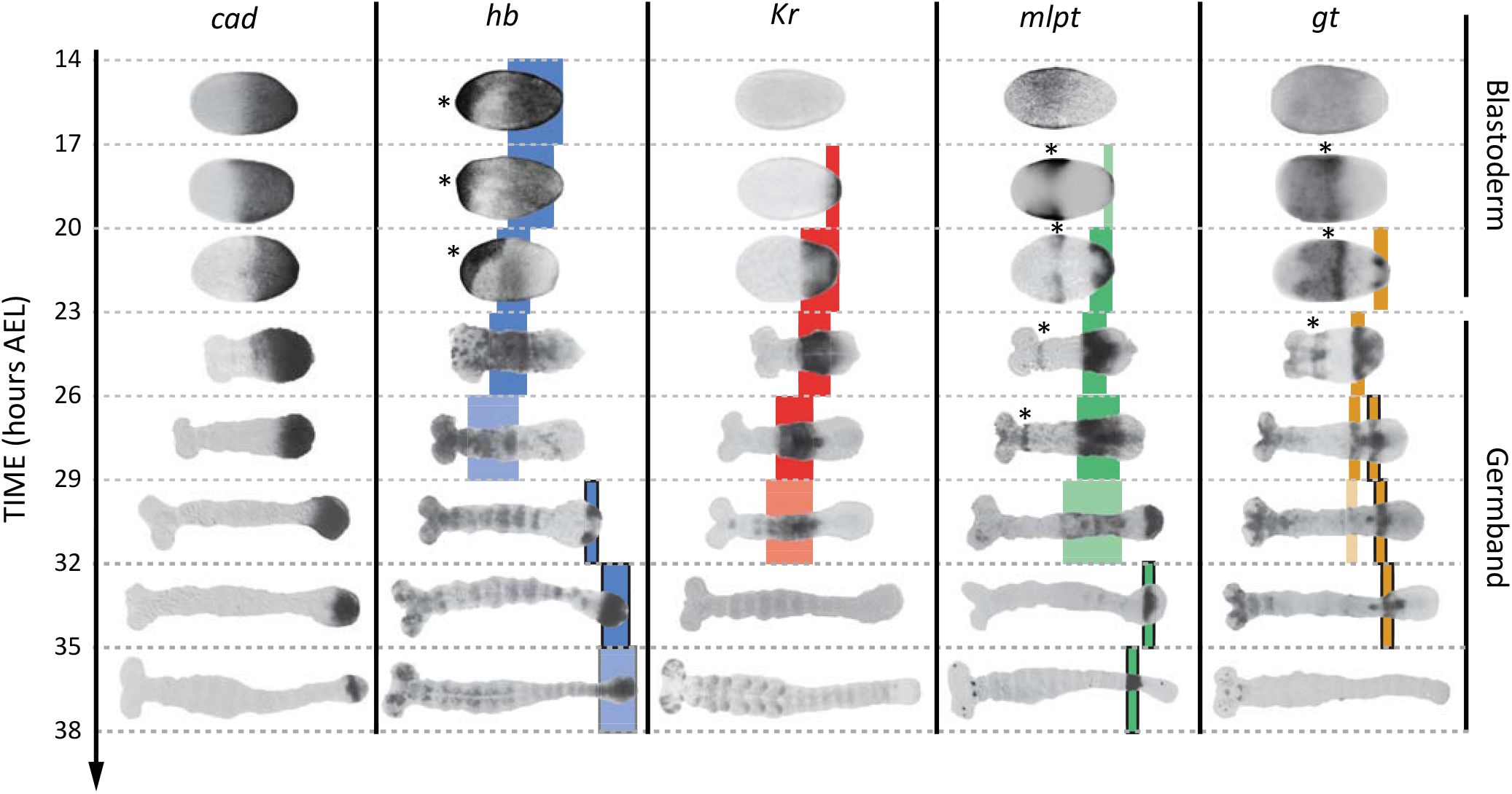
Expression dynamics of *caudal* and gap genes during AP axis specification in *Tribolium*. *caudal* is expressed in a static (i.e. non-retracting) posterior-to-anterior gradient during the blastoderm stage, while is expressed in a retracting posterior-to-anterior gradient during the germband stage. Gap genes are expressed in sequential waves of gene expressions that propagate from posterior to anterior. Expression of different gap genes are tracked with differed colors: blue for *hb*, red for *Kr*, green for *mlpt*, and gold for *gt*. The second trunk domains of *hb, mlpt* and *gt* are outlined in black. Weak expressions are shown in faint colors. Non-trunk and extraembryonic expressions of gap genes (not considered in our analysis) are marked with asterisks. Posterior to the right in all embryos shown.

To determine whether the FF or the SR model is involved in regulating gap genes in *Tribolium*, we sought to re-induce the first gene in the gap gene sequence, namely *hb*, at arbitrary times during AP patterning in the *Tribolium* embryo. To this end, we constructed a transgenic line carrying a *hb* CDS under the control of a heat-shock promoter (hs-hb line; see Materials and Methods). Briefly heat-shocking hs-hb embryos at 26-29 hours After Egg Lay (AEL) indeed resulted in a ubiquitous expression of *hb* that lasted for 6 hours post heat-shock (Figure S3). As a control, we also heat-shocked WT embryos at 26-29 hours AEL, and noticed a normal progression of gap gene expression, albeit with an initial delay of around 9 hours, compared to non-heat-shocked WT embryos (compare Figures 4A and S3 to Figure 3; see also Figure 4C).

**Figure 4.**
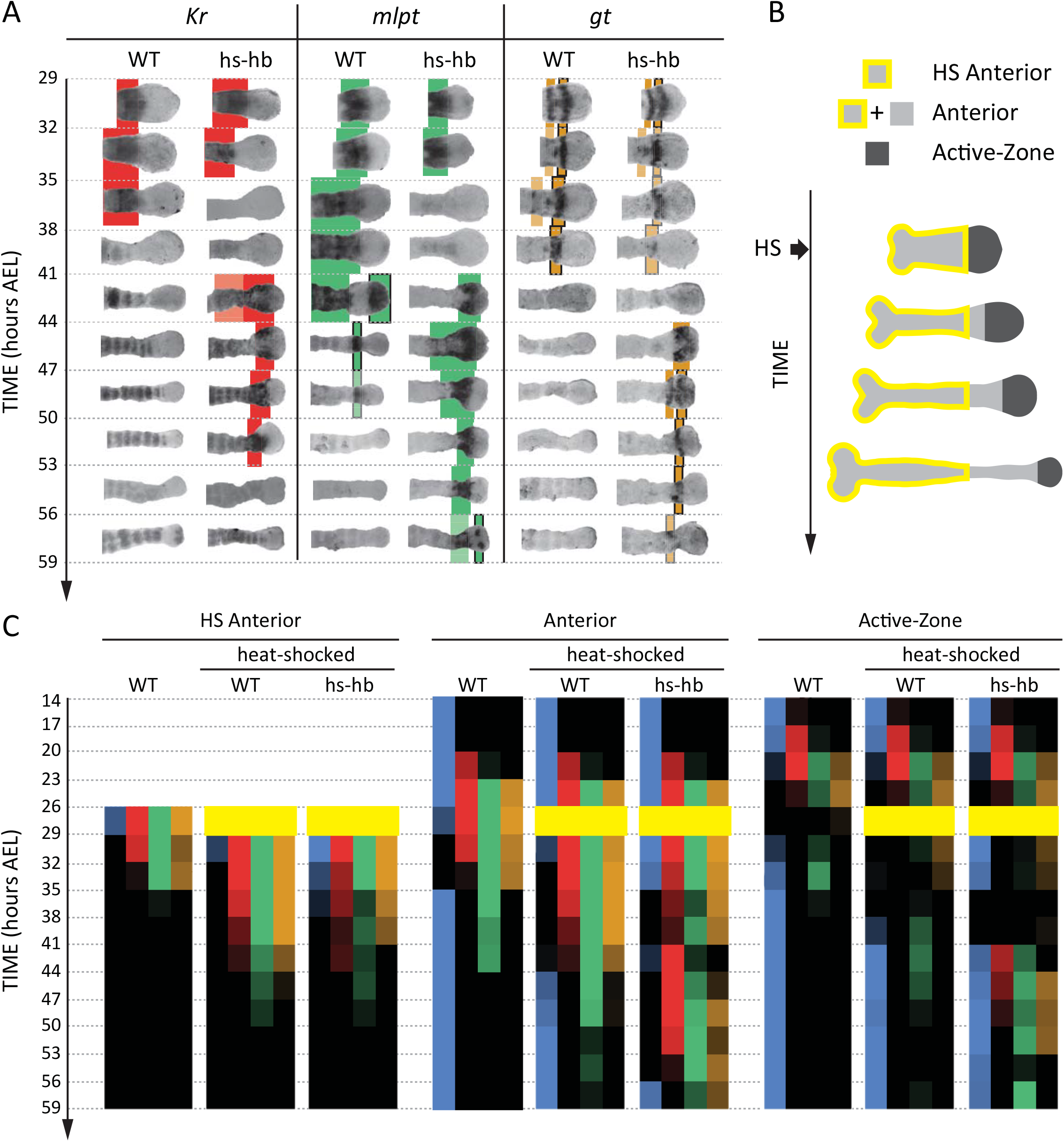
Dual response of *Tribolium* embryo upon re-inducing *hb* expression at 26-29 hours AEL. (**A**) Expression dynamics of gap genes *Kr, mlpt*, and *gt* upon heat-shocking both WT and hs-hb embryos at 2629 hours AEL. Shown are posterior halves of embryos (see Figure S3 for whole embryos). Expression of different gap genes are tracked with differed colors: red for *Kr*, green for *mlpt*, and gold for *gt*. The second trunk domains of *mlpt* and *gt* are outlined in black. Weak expressions are shown in faint colors. Posterior to the right in all embryos shown. (**B**) Dividing heat-shocked *Tribolium* embryos into 3 domains: Active-Zone (caudal-expressing zone at the posterior end of the embryo), Anterior (anterior to the Active-Zone), and HS Anterior (anterior to the Active-Zone at the time of applying heat-shock). (**C**) Quantification (see Methods) of gap gene expressions in WT, heat-shocked WT, and heat-shocked hs-hb embryos (heat-shocks applied at 26-29 hours AEL). Quantifications are carried out separately for HS Anterior, Anterior, and Active-Zone. While heat-shock application resulted in only a temporal delay in gap gene expression in WT, it resulted in dual response for hs-hb embryos: already established gap gene domains are prematurely down-regulated in HS Anterior, while the gap gene sequence is re-activated in Active-Zone. Reinduced gap gene sequence eventually propagates into Anterior. Time-windows where heat-shock is applied are shown in yellow.

Next, we examined and contrasted the expressions of the other gap genes (Kr, *mlpt*, and *gt*) in heat-shocked hs-hb and heat-shocked WT embryos. We observed that cells in hs-hb embryos had two distinct responses depending on whether they are within or anterior to the active-zone. Gene expression domains anterior to the active-zone are pre-maturely repressed, compared to heat-shocked WT. Within the active-zone, the gap gene domains sequence is re-induced, and eventually propagates towards anterior (Figure 4A). Specifically, in heat-shocked hs-hb embryos, at around 35-38 hours AEL, *Kr* expression anterior to the active-zone was down-regulated (Figure 4A). At around 41 hours AEL, *Kr* expression was re-initiated in the active-zone of heat-shocked hs-hb embryos, an effect that is not noticed in heat-shocked WT embryos (Figure 4A). The re-initiated *Kr* expression then propagated towards anterior. A similar effect is observed for the gap gene *mlpt*. At 35 hours AEL, the already established *mlpt* expression at the anterior of hs-hb embryos was down-regulated. By 41 hours AEL, *mlpt* expression was re-established in the active-zone and propagated towards anterior. The second domain of the re-established *mlpt* expression appeared at 56 hours AEL. Similarly, the already formed two domains of *gt* expression were repressed in the anterior and new two domains of expressions were re-established in the posterior of hs-hb embryos that eventually propagated towards anterior (Figure 4A).

We then characterized the response of the *Tribolium* embryo upon *hb* re-induction more quantitatively within and anterior to the active-zone. Since cells within the active-zone progressively move out of this region during axis elongation, we performed our analysis for three regions (Figure 4B): (*i*) the active-zone, (*ii*) anterior to the active-zone at the time of heat-shock application (hereafter called ‘HS Anterior’), and (*iii*) anterior to the active-zone at the time of analysis (hereafter called the ‘Anterior’, which includes HS Anterior and the cells that have moved out of the active-zone since the time of heat-shock application). By examining the distribution of gene activities within these three regions (Active-Zone, HS Anterior, and Anterior) over time, we confirmed the dual response of *Tribolium* embryos upon re-inducing *hb* (Figure 4C). In HS Anterior cells, gap genes are pre-maturely down-regulated; while within the Active-Zone, gap gene sequence is re-induced and eventually propagates towards the Anterior region (as shown in HS Anterior, Active-Zone, and Anterior, respectively in Figure 4C).

It is worth noting here that, in WT, starting from 14 hours AEL, *hb, Kr, mlpt* and *gt* expression domains arise sequentially in the active-zone (Figure 3). By 26 hours AEL, the majority of gap gene expression domains already propagated out of the active-zone towards anterior, and the active-zone becomes virtually void of any gap gene expression. Around 29 hours AEL, the second abdominal domains of *mlpt* and *hb* arise in the active-zone (Figure 3). In Figure 4, we performed our heat-shock experiments within the 26-29 hours AEL time window when the active-zone is void of gap gene expression. The fact that the entire gap gene sequence is re-induced upon re-inducing *hb* in the active-zone at a point in time and space where gap genes are not expressed in WT indicates that gap genes are wired into an ‘aperiodic’ clock that can be reset at any point in time and strongly argues against a threshold-based French Flag model underlying gap gene regulation in *Tribolium* and supports a mechanism based on the Speed Regulation model, as discussed in our earlier theoretical analysis. Furthermore, the dual response of the *Tribolium* embryo to the re-induction of *hb* provides an indirect support for the two-modules GRN realization of the SR model (Figure 1B”). The difference in response between the active-zone and anterior cells can be explained by a difference in the genetic wiring of gap genes in these two regions, encoded by two different GRNs (e.g. dynamic and static modules).

So far, we considered the effects of re-inducing *hb* at 26-29 hours AEL, a time window where the active-zone is void of gap gene expressions. This helped avoiding a possible interference between gap gene expressions already present in the active-zone and the re-induced expressions. To investigate the outcome of re-inducing the gap gene sequence while the original gene expression sequence is unfolding, we preformed our heat-shock experiments also at time windows 23-26 hours AEL (Figure S4) and 20-23 hours AEL (Figure S5). For both time windows, the expression within the active-zone suffered a transient down-regulation before the gap gene sequence is re-induced. This indicates that the gap gene clock is based on a genetic cascade with mutual-repressive links, like the one we used in our theoretical analysis (dynamic module in Figure 1B”).

### The re-induction of gap gene sequence upon re-inducing *hb* is specific to the active-zone

In our earlier theoretical analysis (11), we considered a realization of the SR model that relies on the gradual switching between two genetic modules. An alternative realization would be to jointly regulate the activation and degradation rates of gap genes by a posterior morphogen gradient (Text S1; Movie S9). A major prediction of the module switching model in the case of gap gene regulation in *Tribolium* is that the genetic wiring of gap genes in the presence of the morphogen *cad* (i.e. within the active-zone) is different from their wiring in the absence of *cad* (i.e. anterior to the active-zone). As discussed above, the difference in response to the re-induction of *hb* between the active-zone and the anterior supports the module switching model (and disfavors a degradation rate modulation model; Text S1 and Movie S10). To further test this, we sought to examine if the transient down-regulation and the subsequent re-activation of gap genes is specific to the active-zone, i.e. the region of the embryo expressing *cad*. To this end, we utilized the *axin* (*axn*) RNAi phenotype in *Tribolium*, where the *cad* gradient extends to cover most of the embryo, transforming the embryo into a big active-zone (Figure S6) (11,43). We performed our heat-shock experiments at 20-23 hours AEL time window for WT embryos, hs-hb embryos, embryos laid by WT mothers injected with *axn* dsRNA (*axn* RNAi embryos), and embryos laid by hs-hb mothers injected with *axn* dsRNA (hs-hb; *axn* RNAi embryos). We then analyzed the expression of *mlpt* at consecutive 3-hours time windows starting from 32 hours AEL. As shown earlier, in hs-hb embryos, *mlpt* initially suffered a transient down-regulation. Shortly after, *mlpt* expression is re-established within the active-zone. In *axn* RNAi embryos, *mlpt* expression proceeded as in WT but propagated across the whole embryo and never stabilized, consistent with the fact that the whole embryo transformed into an active-zone (11). In hs-hb; *axn* RNAi embryos, after a transient down-regulation of *mlpt* expression, *mlpt* is activated in the whole embryo, an effect only observed in the active-zone in WT embryos. This supports the hypothesis that the re-activation of gap gene sequence in *Tribolium* upon re-inducing *hb* is specific to the region of the embryo where the posterior morphogen *cad* is expressed.

## Discussion

In this paper, we argued that important classes of GRN realizations of the FF model exhibit dynamic gene expression patterns and are sensitive to morphogen exposure time, at least during an initial transient phase. Nonetheless, such realizations are faithful to the essence of the FF model where morphogen concentration thresholds set the boundaries between different gene expression domains. These thresholds are inscribed either explicitly by tuning the binding affinities of the morphogen gradient to the cis-regulatory elements of downstream genes, implicitly by tuning cross-regulatory strengths between genes or using a combination of both strategies. In contrast, the SR model is essentially threshold-free and drives ever-changing gene expression patterns as long as the morphogen gradient is applied. We also considered a modified version of the FF model (the FFTG model) that, while exhibiting similar spatiotemporal dynamics as SR model, is still threshold-based.

In fact, the SR and the FFTG models are more intimately related than it first appears. Both models can be thought of as composed of an ‘aperiodic’ clock whose speed is regulated by the posterior morphogen (grey in Figures 2B and 2C). The main difference between the two mechanisms is the nature of the clock.

In the SR model, the clock is mediated by the regulatory interactions between the fate-specifying genes themselves (e.g. the dynamic module in Figure 1B”). Each tick of the clock is specified by a different (combination of) fate specifying gene(s). In the suggested molecular realization of the SR model, the speed of the clock can be regulated by the relative proportions of the dynamic and static modules. In the FFTG model, the clock is the Timer Gene, where each tick of the clock is specified by a different concentration of the Timer Gene. Different concentrations of the Timer Gene are translated into different fates by means of a FF model. The speed of the Timer Gene clock is regulated by the activating posterior morphogen (grey in Figure 2C), since the concentration of an activator naturally regulates the transcription rate of its target gene. Although a gene with expression dynamics like that of a Timer Gene has not so far been discovered in insects, its existence is still a viable possibility.

Resetting the clock of the SR model would, then, require resetting the gap gene cascade itself, while resetting the clock of the FFTG model would require resetting the Timer Gene. To investigate whether gap gene regulation in insects are mediate by the SR or the FFTG model, we re-induced the leading gap gene in the gap gene sequence (hb) in the *Tribolium* embryo, which resulted in the re-induction of the gap gene cascade. This provides a strong evidence that gap genes are regulated by a SR model in *Tribolium*.

One limitation of our theoretical analysis and comparison of the SR, FF, and FFTG models is that we used specific GRN realizations for these models, since other GRN realizations could also be possible. However, we used our GRN realizations to illustrate arguments that are general. The GRN realizations of FF (GRN1 and GRN2) and FFTG models are used to show that important classes of FF realizations exhibit similar dynamics to the SR model. The failure of FF GRNs to re-induce the patterning genes sequence upon the re-induction of one of the constituent genes is due to a general feature of the FF model rather than to some modeling specifics of the used GRNs, namely due to that fact that the morphogen gradient is the driving factor behind the sequential activation of genes rather than the cross-regulatory interactions between patterning genes themselves, as the case in the SR model.

In this paper, we used a GRN realization of the SR model based on the gradual switching between two genetic circuits. Another realization of the SR model would be to regulate both the activation and decay rates of the constituent genes of a genetic cascade by the morphogen gradient. However, the dual response of *Tribolium* embryos upon re-inducing *hb* supports the two-module realization (compare Movies S7 and S10). Yet another possible GRN realization for the SR module is the composite AC/DC mode of the AC-DC motif (44). The AC-DC GRN can operate either as a multi-stable circuit (the DC mode) or as a clock (the AC mode), depending on the concentration of a morphogen gradient. At certain ranges of the morphogen gradient, both modes could co-exist, resulting in the fine-tuning of the speed of the clock. Hence, the AC-DC GRN has a somewhat similar logic as the two-modules GRNs; however, it uses one gene circuit that can work as both dynamic or static modules depending on an external factor rather than explicitly using two genetic modules as in the two-modules GRN. However, it is not clear if the AC-DC GRN would be able to mediate a morphogen-mediated transition from the composite AC-DC mode to a pure DC mode to fully realize wavefront-based patterning. In addition, while the AC-CD GRN is economical as it uses one module to achieve both AC and DC behaviors, it lacks the flexibility of the two-modules model, since different realizations of the dynamic and static modules in the two-modules model could be employed to mediate a wide range of final spatial patterns without the constraint of using one module to mediate two functions.

In summary, in this paper we showed that the gene expression dynamics driven by a French Flag mechanism can be indistinguishable from those driven by a threshold-free mechanism. We then suggested a test to differentiate between the two cases and carried it out experimentally in the beetle *Tribolium castaneum*. The test confirmed that the AP fate specification in *Tribolium* (and possibly all insects) is based on the speed regulation of an aperiodic clock of gap genes in a threshold-free fashion.

**Figure 5.**
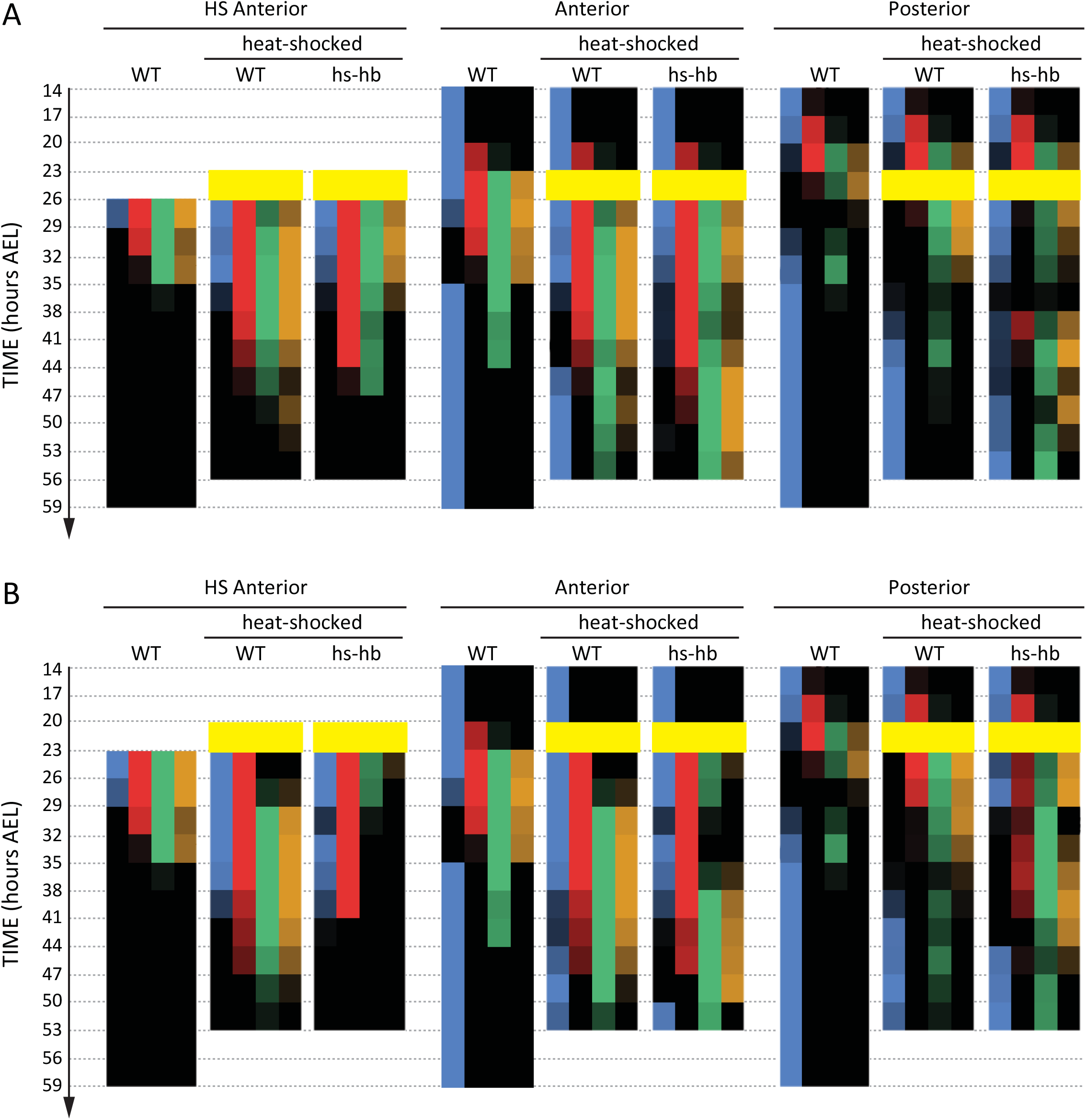
Dual response of *Tribolium* embryo upon re-inducing *hb* expression at 23-26 and 20-23 hours AEL. Quantification (see Methods) of gap gene expressions in WT, heat-shocked WT, and heat-shocked hs-hb embryos. Heat-shocks are applied at 23-26 hours AEL (**A**) and 20-23 hours AEL (**B**). Quantifications are carried out separately for HS Anterior, Anterior, and Active-Zone. While heat-shock application resulted in only a temporal delay in gap gene expression in WT, it resulted in dual response for hs-hb embryos: already established gap gene domains are pre-maturely down-regulated in HS Anterior, while the gap gene sequence is re-activated in Active-Zone. Re-induced gap gene sequence eventually propagates into Anterior. Time-windows where heat-shock is applied are shown in yellow.

**Figure 6.**
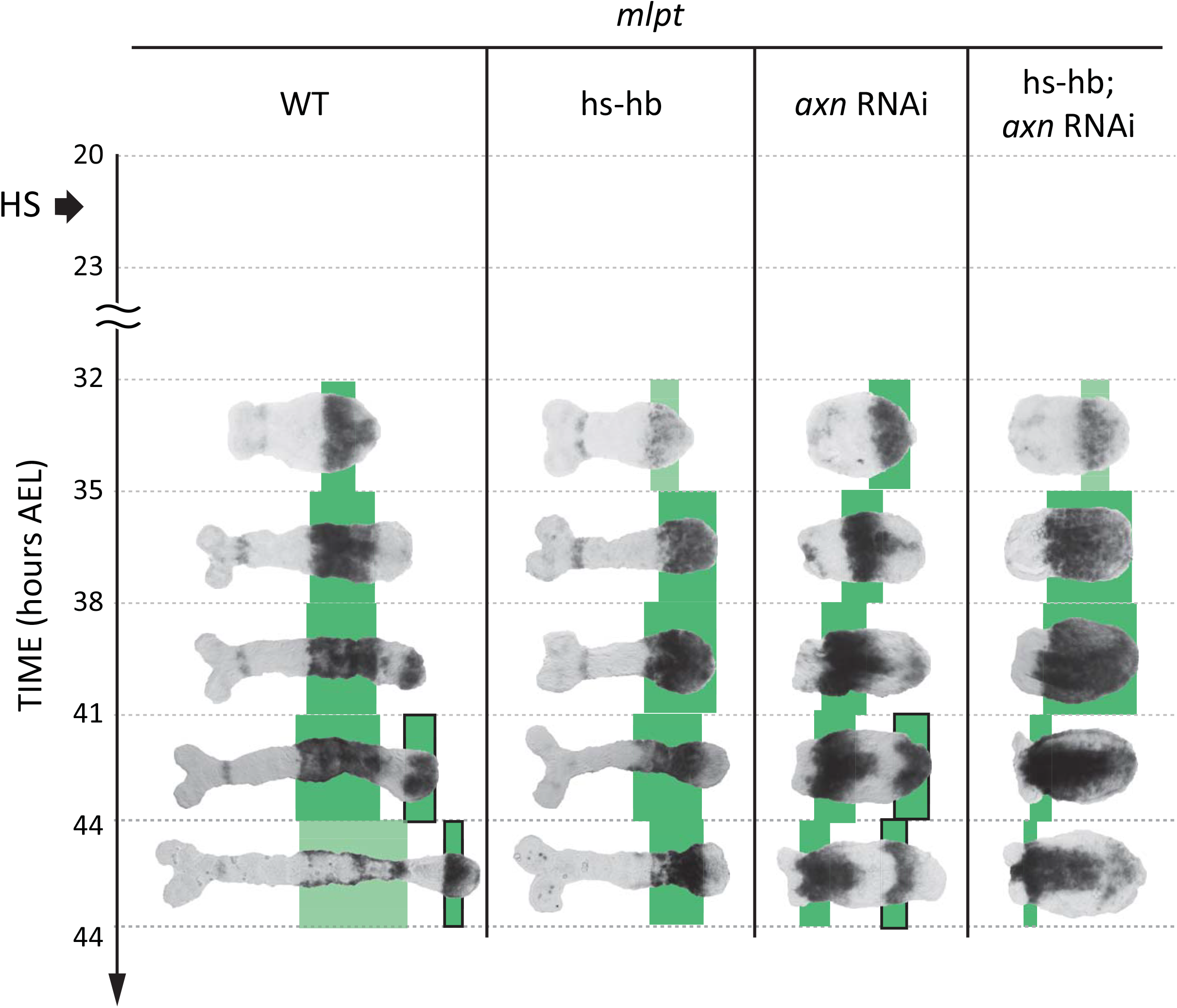
Gap gene sequence re-activation upon re-inducing *hb* is specific to the active-zone. *mlpt* expression is re-activated only in the active-zone (posterior cad-expressing region) upon heat-shocking hs-hb embryos (compare WT and hs-hb embryos). Knocking-down *axn* completely posteriorized *Tribolium* embryos (*axn* RNAi), such that nearly the entire embryo becomes a big active-zone (as evident from *cad* expression; see Figure S6), where *mlpt* expression is very dynamic and propagates across the entire embryo. Heat-shocking hs-hb embryos whose mothers had been injected with *axn* dsRNA (hs-hb; *axn* RNAi) re-activates *mlpt* expression in the whole embryo, supporting the hypothesis that gap gene reactivation in specific to the cad-expressing domain (the active-zone). *mlpt* expresssion is tracked in green. The second trunk domain of *mlpt* is outlined in black. Weak expressions are shown in faint colors. Posterior to the right in all embryos shown.

---

**Figure S1.**
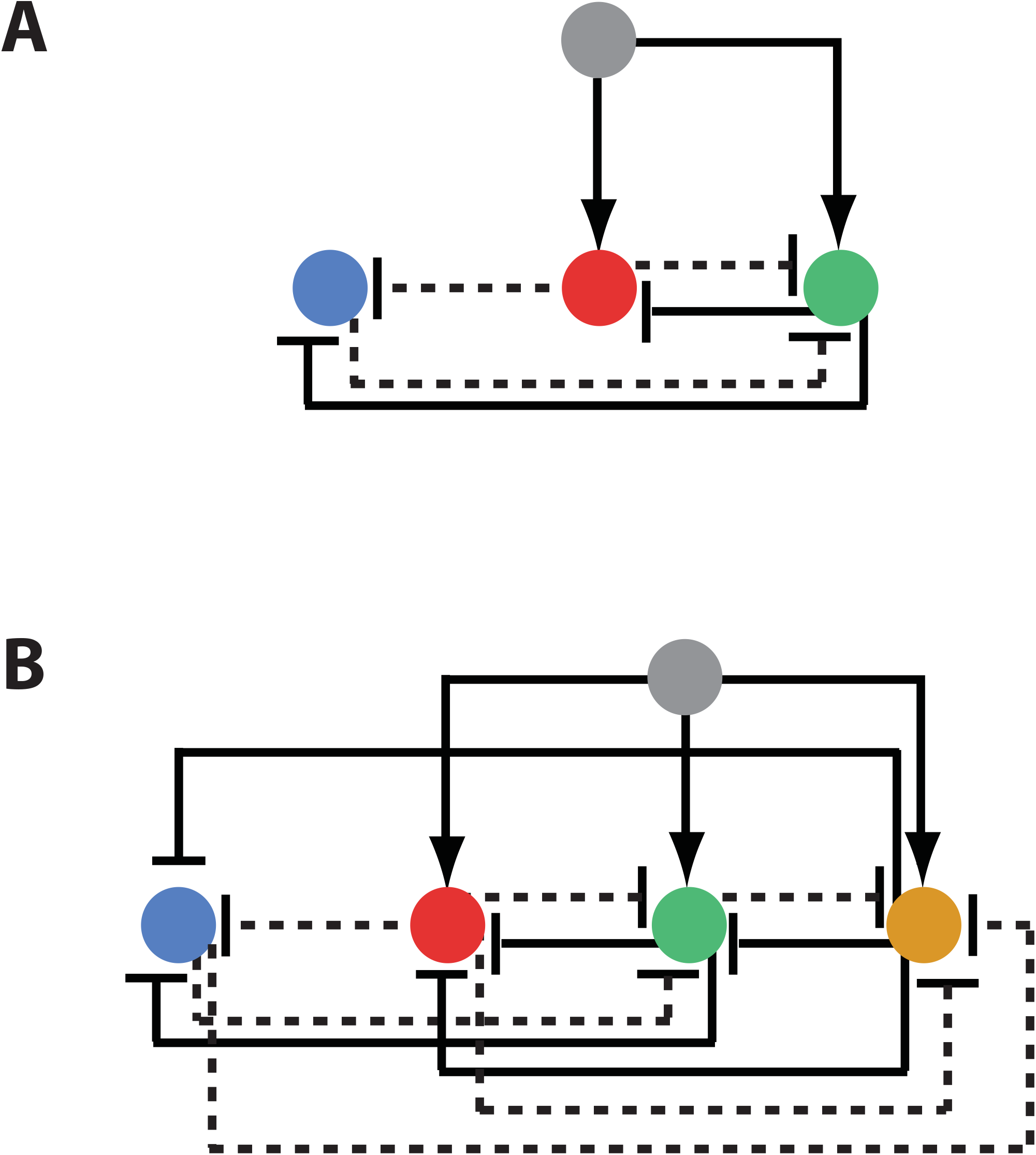
French Flag GRN 2 (or AC-DC network motif). (**A**) Three-genes French Flag GRN 2. (**B**) Four-genes French Flag GRN 2. Grey gene: morphogen gradient. Dashed lines signify weak activation/repressions.

**Figure S2.**
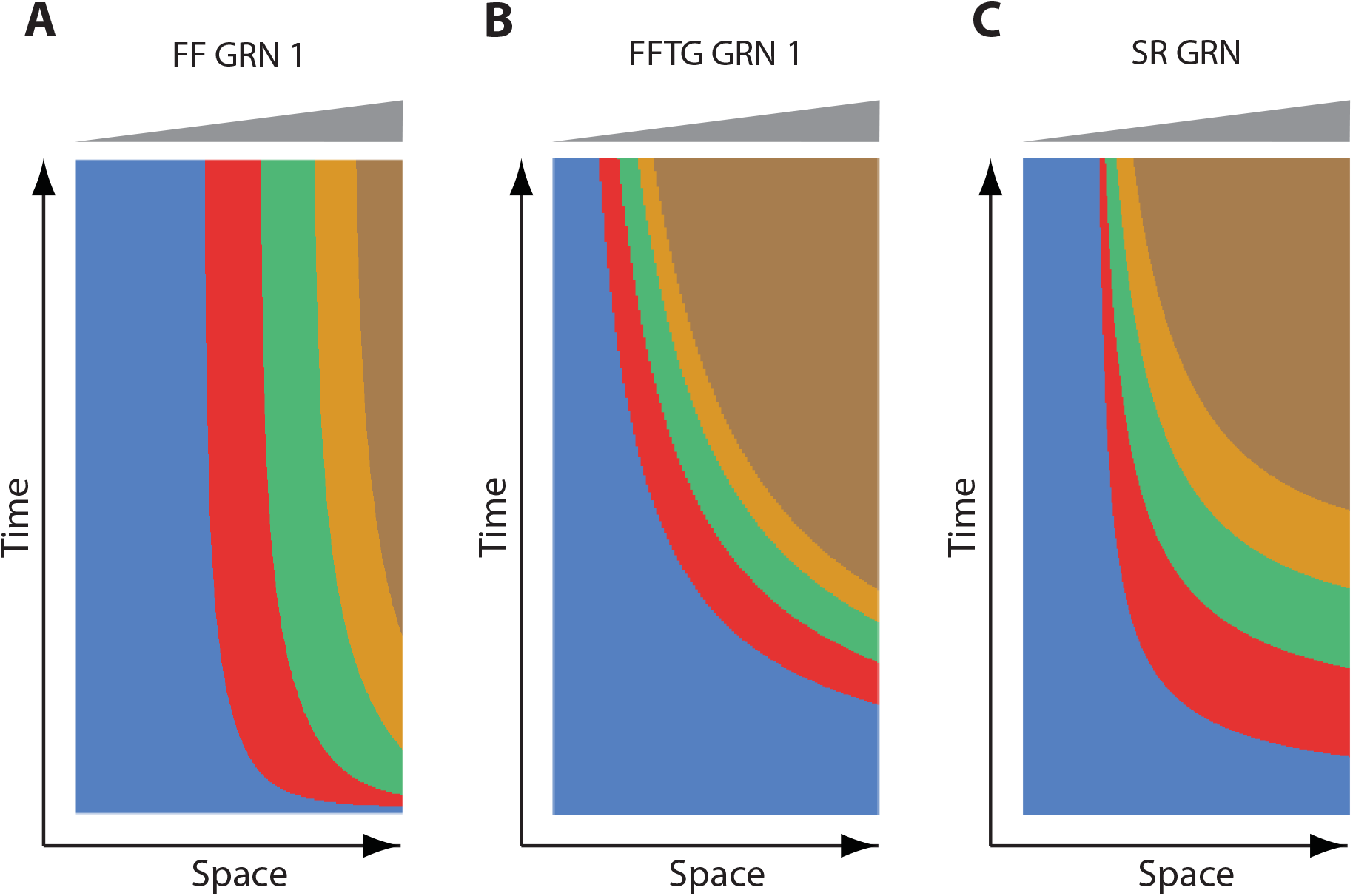
Steady state behavior of French Flag GRN 1, French Flag with Timer Gene GRN, and Speed Regulation GRN. Shown are kymographs of computer simulations of French Flag GRN 1 (FF GRN 1), French Flag with Timer Gene GRN (FFTG), and Speed Regulation (SR) GRN. For FF GRN 1, genes are expressed initially in sequential waves, but eventually reach steady state. For FFTG and SR GRNs, genes are expressed in sequential waves that keep propagating and shrinking as long as the morphogen gradient (shown in grey) is applied.

**Figure S3.**
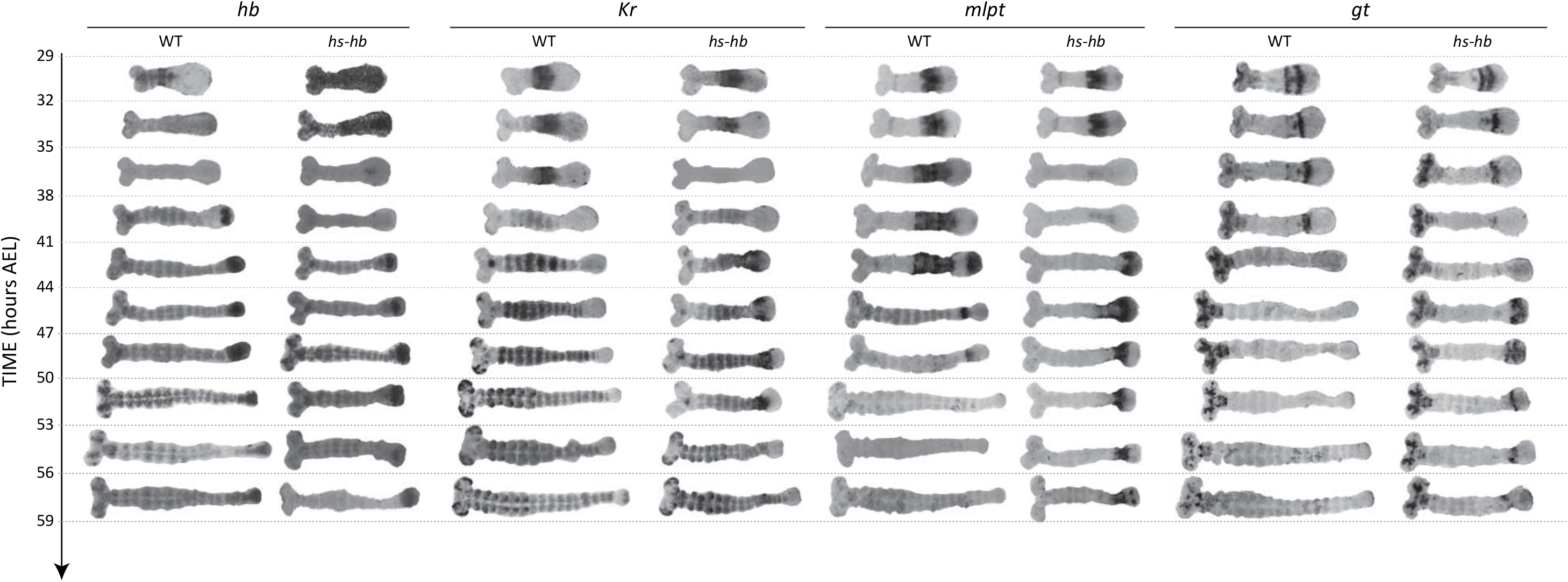
Dual response of *Tribolium* embryo upon re-inducing *hb* expression at 26-29 hours AEL. Expression dynamics of gap genes *hb, Kr, mlpt*, and *gt* upon heat-shocking both WT and hs-hb embryos at 26-29 hours AEL. Posterior to the right.

**Figure S4.**
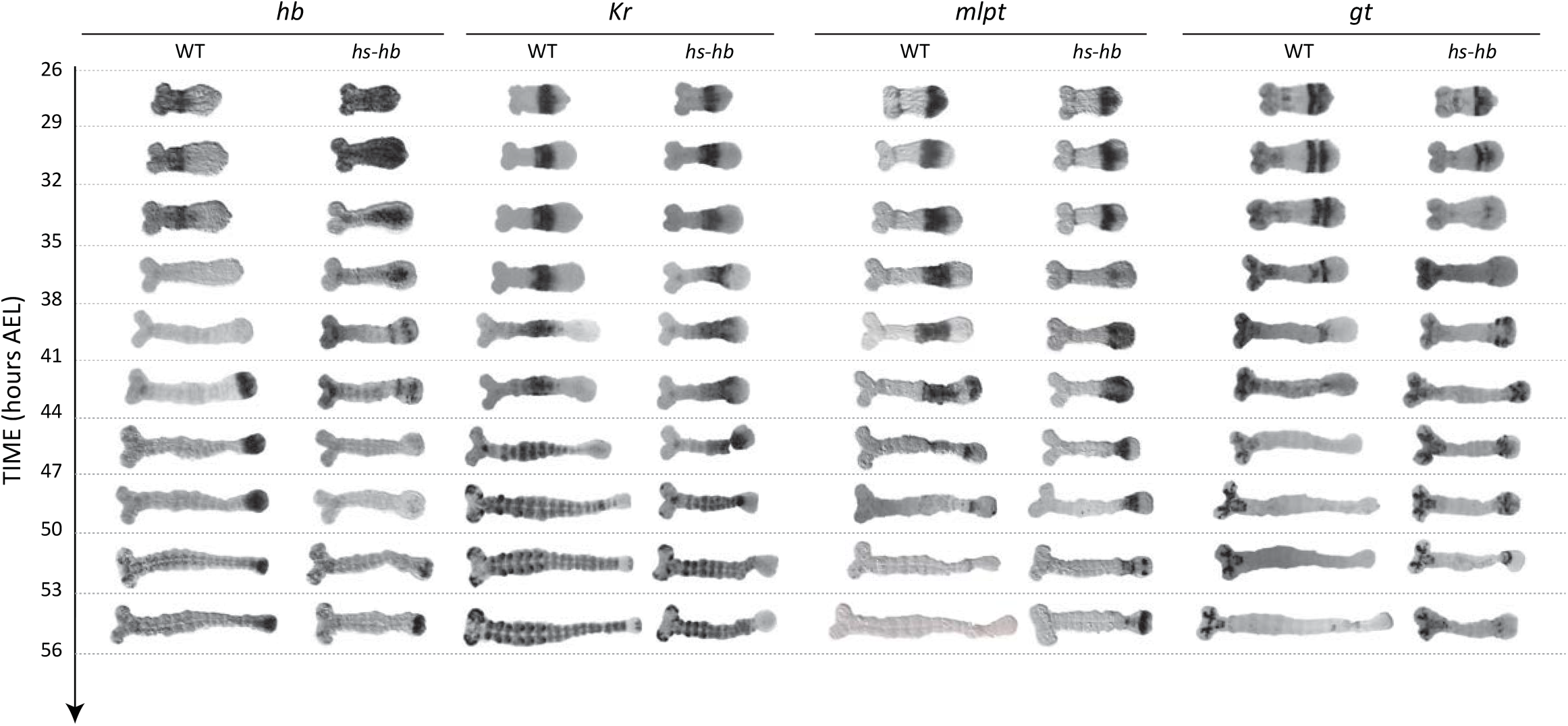
Dual response of *Tribolium* embryo upon re-inducing *hb* expression at 23-26 hours AEL. Expression dynamics of gap genes *hb, Kr, mlpt*, and *gt* upon heat-shocking both WT and hs-hb embryos at 23-26 hours AEL. Posterior to the right.

**Figure S5.**
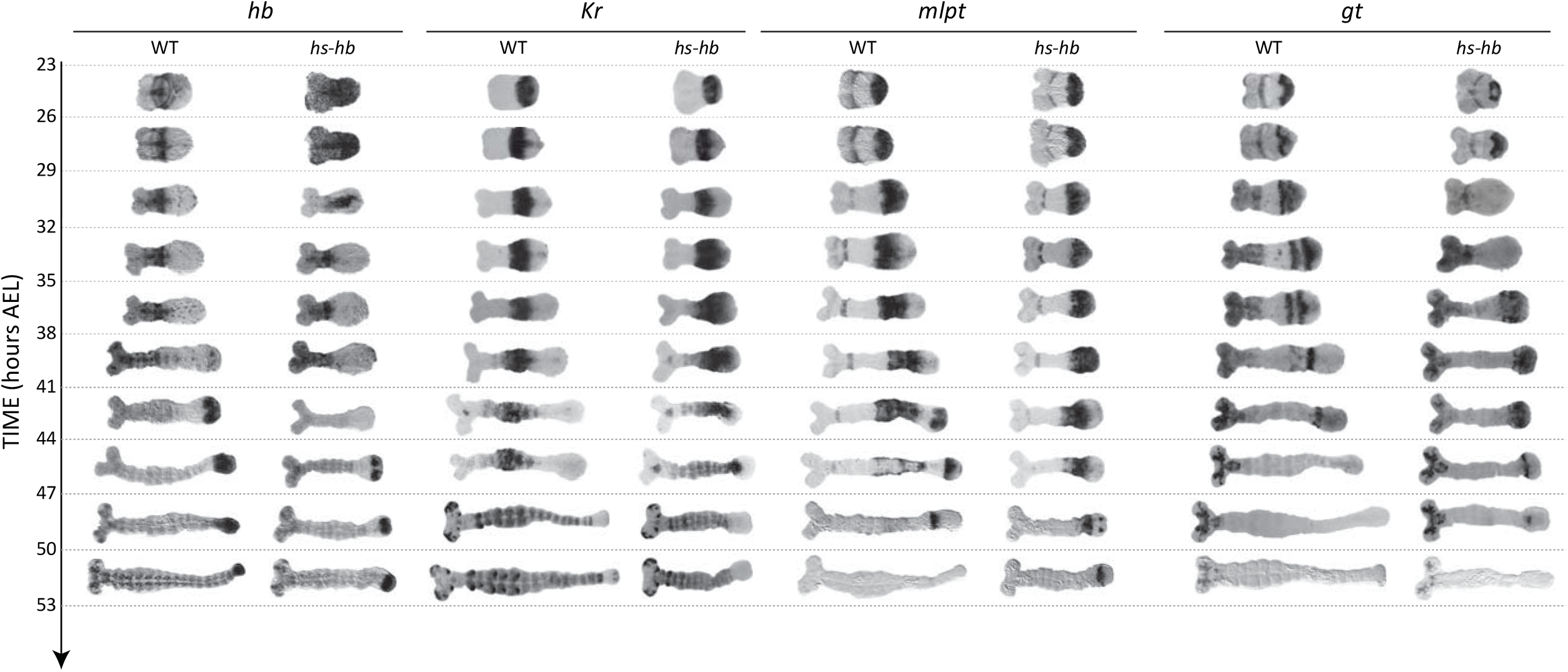
Dual response of *Tribolium* embryo upon re-inducing *hb* expression at 20-23 hours AEL. Expression dynamics of gap genes *hb, Kr, mlpt*, and *gt* upon heat-shocking both WT and hs-hb embryos at 20-23 hours AEL. Posterior to the right.

**Figure S6.**
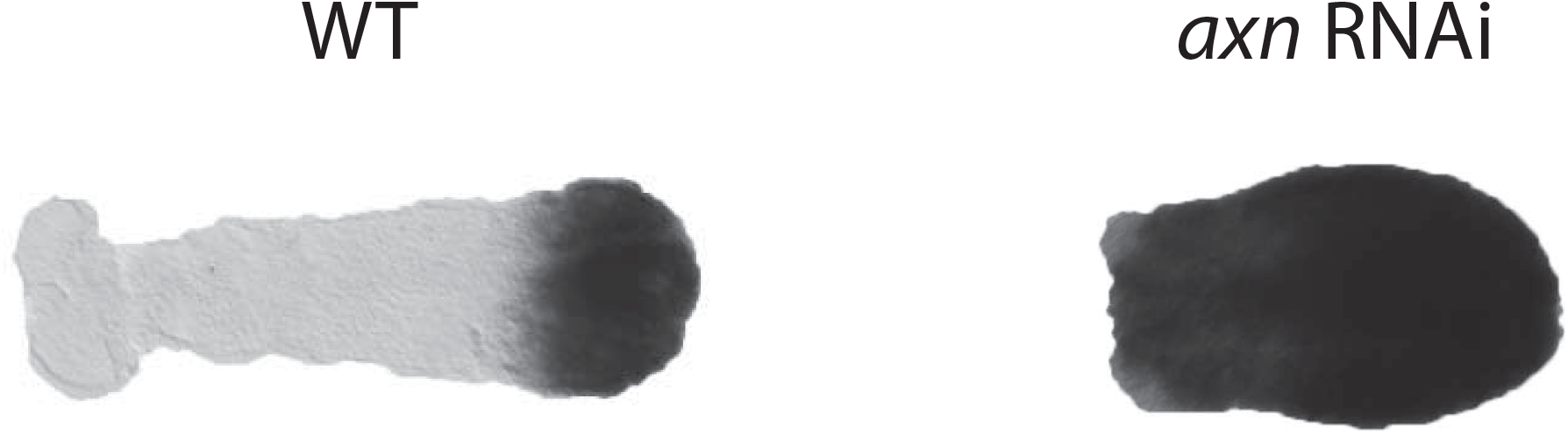
Knocking-down *axn* by RNAi posteriorizes *Tribolium* embryos. Shown is *cad* expression in WT and *axn* RNAi embryos. *cad* expression is restricted to the posterior end in WT, while *cad* expression extends to cover most of *axn* RNAi embryos upon loss of this anterior repressor of Wnt activity which activates *cad* (43).

**Movie S1. Computer simulation of French Flag GRN 1.** A computer simulation of a 5-genes French Flag GRN 1 (a 5-genes version of Figure 1A”). Genes are initially expressed in sequential waves, but eventually reaches a steady state. Patterning genes are shown in blue, red, green, gold, and brown. Morphogen gradient is shown in grey. Horizontal axis is space and vertical axis is gene expression concentration.

**Movie S2. Computer simulation of French Flag GRN 2 (AC-DC circuit motif).** A computer simulation of a 3-genes French Flag GRN 2 (see Figure S1 A). Genes are initially expressed in sequential waves, but eventually reaches a steady state. Patterning genes are shown in blue, red, and green. Morphogen gradient is shown in grey. Horizontal axis is space and vertical axis is gene expression concentration.

**Movie S3. Computer simulation of Speed Regulation GRN.** A computer simulation of a 5-genes Speed Regulation GRN (a 5-genes version of Figure 1B”). Genes are expressed in sequential waves that never stabilizes (except for a small region at the low end of the morphogen). Patterning genes are shown in blue, red, green, gold, and brown. Morphogen gradient is shown in grey. Horizontal axis is space and vertical axis is gene expression concentration.

**Movie S4. Computer simulation of Speed Regulation GRN driven by a decaying morphogen gradient.** A computer simulation of a 5-genes Speed Regulation GRN (a 5-genes version of Figure 1B”) driven by a continuously decaying morphogen gradient (grey). Genes are expressed in sequential waves that gradually stabilizes due to the morphogen gradient decay. Patterning genes are shown in blue, red, green, gold, and brown. Morphogen gradient is shown in grey. Horizontal axis is space and vertical axis is gene expression concentration.

**Movie S5. Speed Regulation GRN patterning the Anterior-Posterior axis of short-germ, intermediate-germ and long-germ insects.** A computer simulation of fate specification in insects with different germ types using a 5-genes Speed Regulation GRN. (**A**) In short-germ insects, the posterior morphogen (grey) continuously retracts towards posterior with axis elongation. (**B**) In intermediate-germ insects, the posterior morphogen is initially expressed in a static gradient that eventually retracts towards posterior with axis elongation. (**C**) In long-germ insects, the posterior morphogen is expressed in a static gradient throughout the patterning process. Patterning genes are shown in blue, red, green, gold, and brown. Posterior morphogen gradient is shown in grey. Horizontal axis represents the Anterior-Posterior axis. Posterior to the right. Vertical axis is gene expression concentration.

**Movie S6. French Flag with a Timer Gene GRN patterning the Anterior-Posterior axis of short-germ, intermediate-germ and long-germ insects.** A computer simulation of fate specification in insects with different germ types using a 5-genes French Flag with a Timer Gene GRN. (**A**) In short-germ insects, the posterior morphogen (grey) continuously retracts towards posterior with axis elongation. (**B**) In intermediate-germ insects, the posterior morphogen is initially expressed in a static gradient that eventually retracts towards posterior with axis elongation. (**C**) In long-germ insects, the posterior morphogen is expressed in a static gradient throughout the patterning process. Patterning genes are shown in blue, red, green, gold, and brown. Posterior morphogen gradient is shown in grey. Timer Gene is shown in black. Horizontal axis represents the Anterior-Posterior axis. Posterior to the right. Vertical axis is gene expression concentration.

**Movie S7. Re-inducing the leading gene in the Speed Regulation GRN during intermediate-germ patterning simulation.** Re-inducing the leading gene (blue) in the Speed Regulation GRN during a simulation of intermediate-germ patterning results in dual response: already established genes in the anterior are down-regulated, while the sequential activation of genes is set within the expression domain of the posterior morphogen (grey). Patterning genes are shown in blue, red, green, gold, and brown. Posterior morphogen gradient is shown in grey. Horizontal axis represents the Anterior-Posterior axis. Posterior to the right. Vertical axis is gene expression concentration.

**Movie S8. Re-inducing the leading gene in the French Flag with a Timer Gene GRN during intermediate-germ patterning simulation.** Re-inducing the leading gene (blue) in the French Flag with a Timer Gene GRN during a simulation of intermediate-germ patterning results in a transient dominance of the leading gene, but eventual formation of the normal gene expression pattern. Patterning genes are shown in blue, red, green, gold, and brown. Posterior morphogen gradient is shown in grey. Horizontal axis represents the Anterior-Posterior axis. Posterior to the right. Vertical axis is gene expression concentration.

**Movie S9. Simulation of GRN realizing the Speed Regulation model by jointly modulating gene activity and gene products decay rates.** A computer simulation of the Speed Regulation model realized by jointly modulating gene activity and gene products decay rates (described in Text S1) applied to the problem of patterning the anterior-posterior axis of an intermediate-germ insect. Patterning genes are shown in blue, red, green, gold, and brown. Posterior morphogen gradient is shown in grey. Horizontal axis represents the Anterior-Posterior axis. Posterior to the right. Vertical axis is gene expression concentration.

**Movie S10. Re-inducing the leading gene in the Speed Regulation GRN realization by jointly modulating gene activity and gene products decay rates during intermediate-germ patterning simulation.** Re-inducing the leading gene (blue) in the Speed Regulation GRN realization by jointly modulating gene activity and gene products decay rates during a simulation of intermediate-germ patterning results resetting the sequential activation of patterning genes within the expression of the posterior morphogen (grey) but not leaves the already established gene expression in the anterior intact. Patterning genes are shown in blue, red, green, gold, and brown. Posterior morphogen gradient is shown in grey. Horizontal axis represents the Anterior-Posterior axis. Posterior to the right. Vertical axis is gene expression concentration.

## Materials and Methods

### in situ hybridization, RNAi, and imaging of fixed embryos

*In situ* hybridization was performed using DIG-labeled RNA probes and anti-DIG::AP antibody (Roche). Signal was developed using NBT/BCIP (BM Purple, Roche). All expression analyses were performed using embryos from uninjected females or females injected with double-stranded RNA (ds RNA) of gene of interest. dsRNA was synthesized using the T7 megascript kit (Ambion) and mixed with injection buffer (5 mM KCl, 0.1 mM KPO4, pH 6.8) before injection. Used dsRNA concentrations for *axn* RNAi: _ ng/μl Blastoderm stage embryos were imaged with a Nikon Digital DXM 1200F camera on an Olympus BX50 microscope using Nikon ACT-1 version 2.62 software. Germband stage embryos were imaged ProgRes CFcool camera on Zeiss Axio Scope.A1 microscope using ProgRes CapturePro image acquisition software. Brightness and contrast of all images were adjusted and placed on a white background using Adobe Photoshop.

### Overexpression construct

A 1757 bp fragment of the *Tc-hb* mRNA (cDNA22, see (Wolff et al., 1995) was amplified using PCR primers Tc-hb_left 5’-CGTCTAGAGCAAAAATTTCGAACAGTCG-3’ and Tc-hb_right 5’-CCGCTCGAGTCCAACCCGTACATCTCCAT-3’ which were designed to include restriction sites XbaI and XhoI, respectively. This *Tc-hb* fragment contains the complete coding region and partial 5’UTR and 3’UTR sequences and was cloned between a *Tc’hsp68* promotor and 3’UTR sequences via the XbaI-XhoI sites in plasmid pSLfa[Tc’hsp5’-dsRedEx-3’UTR; 5’3’UTR]fa (Schinko and Bucher, unpublished). From this plasmid, an AscI-FseI fragment including the hs-hb cassette was subcloned as AscI-FseI fragment into the piggyBac transformation vector pBac[3xP3-gTcv] (Schinko unpublished), resulting in pBac[3xP3-v; Tc’hsp68-Tc’hb-Tc’hsp68 3’UTR] (see supplementary Fig. S1). This vector uses the *Tribolium vermilion* gene as transformation marker (Lorenzen et al., 2002).

### Generation of transgenic beetles

Plasmid DNAs were isolated using the Quiagen plasmid Midi Kit, and germline transformation was performed as described in (Berghammer et al., 1999; Berghammer et al., 2009). For hs-hb, in one experimental series 408 embryos were injected of which 22% hatched. 44 inter se crosses were set up, from which 10 transgenic strains could be generated. In another experimental series, 210 embryos were injected with a hatchrate of 56%. 59 inter se single matings resulted in 5 transgenic lines.

10 of these hs-hb lines were tested for heatschock phenotypes. Phenotype strength was measured by determining the proportion of larvae which (1) developed homeotic transformations, and (2) which displayed, in addition to the homeotic transformations, additional trunk segments. Two out of those ten lines, which seemed most effective in generating heat-shock phenotypes, were further studied and the strain hs-hb 2 was used to generate most data in this paper.

### Non-heat-shocked egg collections

Three hours developmental windows were generated by incubating three-hours egg collections at 23–24°C for the desired length of time before fixation. Beetles were reared in whole-wheat flour supplemented with 5% dried yeast.

### Heat-shocked egg collections

Three hours developmental windows were generated by incubating three-hours egg collections at 23–24°C for the desired length of time. Egg collections are then heat-shocked in a water bath at 48°C for 10 minutes and then re-incubated at 23–24°C for the desired length of time before fixation.

### Quantification of gene expressions in HS Anterior, Anterior, and Active-Zone

Gene expression quantifications (Figure 4C and Figure 5) were created by counting percentages of embryos that have detectable expression in the three regions: HS Anterior, Anterior, and Active-Zone. Dividing an embryo into HS Anterior, Anterior, and Active-Zone is carried out as shown in Figure 4B.

### Computational Modeling

See Text S1.

